# Distinct spatiotemporal contribution of morphogenetic events and mechanical tissue coupling during Xenopus neural tube closure

**DOI:** 10.1101/2021.06.01.446407

**Authors:** Neophytos Christodoulou, Paris A. Skourides

**Affiliations:** Department of Biological Sciences, University of Cyprus, P.O. Box 20537, 2109 Nicosia, Cyprus

## Abstract

Neural tube closure (NTC) is a fundamental process during vertebrate embryonic development and is indispensable for the formation of the central nervous system. Here, using *Xenopus laevis* embryos, live imaging, single cell tracking, optogenetics and loss of function experiments we examine the contribution of convergent extension (CE) and apical constriction (AC) and we define the role of the surface ectoderm (SE) during NTC. We show that NTC is a two-stage process and that CE and AC do not overlap temporally while their spatial activity is distinct. PCP driven CE is restricted to the caudal part of the neural plate (NP) and takes place during the first stage. CE is essential for correct positioning of the NP rostral most region in the midline of the dorsoventral axis. AC occurs after CE throughout the NP and is the sole contributor of anterior NTC. We go on to show that the SE is mechanically coupled with the NP providing resistive forces during NTC. Its movement towards the midline is passive and driven by forces generated through NP morphogenesis. Last, we show that increase of SE resistive forces is detrimental for NP morphogenesis, showing that correct SE development is permissive for NTC.

## Introduction

One of the most important events during vertebrate embryogenesis is the formation of the central nervous system. The first critical event during central nervous system development is the formation of the neural tube (NT). The NT emerges from the dorsal ectoderm and is the precursor of the brain and spinal cord[1]. Defective NT formation leads to neural tube defects(NTDs) which are some of the most common human birth defects, occurring in 0.5-2 per 1000 births[2]. Thus, understanding the morphogenetic events driving NT formation will provide a better understanding of human NTDs and facilitate their prevention.

The transformation of the flat neuroepithelium into the NT during primary neurulation is facilitated by intrinsic forces generated by morphogenetic events within the neural plate (NP). Studies in different vertebrate models identified two distinct morphogenetic processes controlling the active remodelling of the neuroepithelium during neural tube closure (NTC). Specifically, convergent extension (CE)[3-5] and apical constriction (AC) [6-8] are the two main morphogenetic movements taking place within the NP as it transforms into the NT.

CE is a morphogenetic process driven by polarized cell intercalation, leading to tissue lengthening and narrowing[9, 10]. Studies in mouse, frog and chicken embryos demonstrated that the planar cell polarity (PCP) pathway governs CE during NTC, as abrogation of PCP signalling leads to impaired CE in this tissue [4, 5, 10-12]. Specifically, PCP controls the polarised activation of actomyosin[4, 13], which is essential for polarised junction remodelling and subsequent neighbour exchanges[14-17].

AC is a morphogenetic process driving the conversion of columnar cells to wedge shaped as a result of apical surface area reduction[18]. We and others have previously shown that during Xenopus NTC AC is controlled by calcium transients and asynchronous medial actomyosin contraction events[8, 19]. AC is responsible for the generation of forces resulting in epithelial folding, driving the bending of the neuroepithelium in vertebrates[20-22]. Studies in mouse and frog embryos highlight the importance of AC during primary neurulation, since genetic ablation of genes regulating AC results in NTDs[6, 7, 23, 24].

Interestingly, while defects in both CE and AC result in NTDs, failure of CE is correlated with spinal NTDs whereas failure of AC is correlated with rostral NTDs[5-7, 12, 25]. The latter suggests that while these two morphogenetic processes are both indispensable for NTC, they have discrete spatial and temporal contribution during NT morphogenesis. At the moment the precise spatiotemporal contribution of these processes towards NTC is not clear.

A number of studies have suggested that in addition to NP intrinsic morphogenesis, extrinsic forces generated by neighbouring tissues contribute to NP morphogenesis [10, 26, 27]. Specifically, it has been proposed that the surface ectoderm (SE) generates a pushing forces for driving the medial movement of the neural folds. In support of this, a study suggested that active cell movement in the deep SE generates a pushing force during NTC[28]. However, NP explants lacking the SE could still form a NT[29]. Additionally the tension within the SE is not anisotropic, which is contradictory with the notion that the SE generates a medially directed pushing force[27], necessary for NTC.

Here we aim to understand the spatiotemporal contribution of the morphogenetic process within the NP and delineate the role of the SE in NTC. Using live imaging and single cell tracking we show that the caudal and rostral regions of the NP display distinct behaviours. This distinct behaviour is the result of differential PCP activity within the two regions. PCP at the caudal region drives CE of the tissue which pushes the rostral regions of the NP forward, and positioning the anterior most region in the midline of the dorsoventral axis. Moreover, we show that AC follows CE throughout the NP and is the sole contributor of anterior NTC. Subsequently we show that the SE doesn’t display active migratory behaviour during NTC, and its medial movement is dependent on NP morphogenesis. Last, we show that a balance between the forces generated by NP morphogenesis and SE tension, is required for NTC.

## Results

### Macroscopic analysis of neural tube closure reveals differential behaviour of caudal and rostral domains

NTC is a process controlled by distinct morphogenetic events within the NP. We and others have previously focused on how single cell behaviour contributes to NTC [8, 17]. However, our understanding of how these events are translated into collective tissue behaviour is lacking. Thus, we decided to map collective tissue movements during NTC with single cell resolution. To achieve this, we generated high temporal resolution time lapse recordings of Xenopus embryos during NTC. Thereafter, we used a low magnification objective in order to utilize the wide field of view and follow the behaviour of the neuroepithelium over a time period of 2-4h (**Fig. 1A**). We performed microinjections of histone-RFP mRNA at animal site of the two blastomeres of 2-cell stage embryos in order to use the nuclear signal to track single cells. The embryos were allowed to develop to stage 12.5-13 and subsequently imaged. We collected 50-60 Z-stacks of 8um every 3 minutes to image the dorsal half of the embryo. After the generation of time lapse recordings, we tracked single cells (**Fig. 1B-C, Movie 1**). Overall, we tracked 816 cells (468 neuroepithelial and 348 SE cells). This revealed that neuroepithelial cells display district motion patterns along the anteroposterior (AP) axis of the embryo. Specifically, the posterior NP (spinal cord-hindbrain region) lengthens and shortens, while the anterior NP’s (midbrain-forebrain region) shape remains the same over a time period of 2 hours (**Fig 1C-E, Movie 1**). Importantly, examination of neuroepithelial cell movement revealed distinct patterns of movement at the anterior and posterior part of the tissue. Specifically, the anterior NP displays a linear movement polarized towards the embryo’s anterior ventral side, while the posterior NP moves towards the midline (**Fig. 1F-G**). At the same time cell proliferation is uniform throughout the NP (**Fig. 1H, Movie 2**), ruling out any contribution of cell proliferation in the differential behaviour or the tissue along the A/P axis. Overall, these data show that the caudal and rostral regions of the NP behave differently during NTC, suggesting that distinct morphogenetic processes occur along the AP axis of the tissue. Lengthening and narrowing of tissues is known to be driven by PCP mediated intercalative behaviour during CE[4, 13, 30]. Thus, together these data suggest that anterior and posterior NP have different intrinsic patterns of morphogenesis due to the presence of PCP mediated intercalative behaviour only at the posterior part of the tissue.

**Figure 1.**
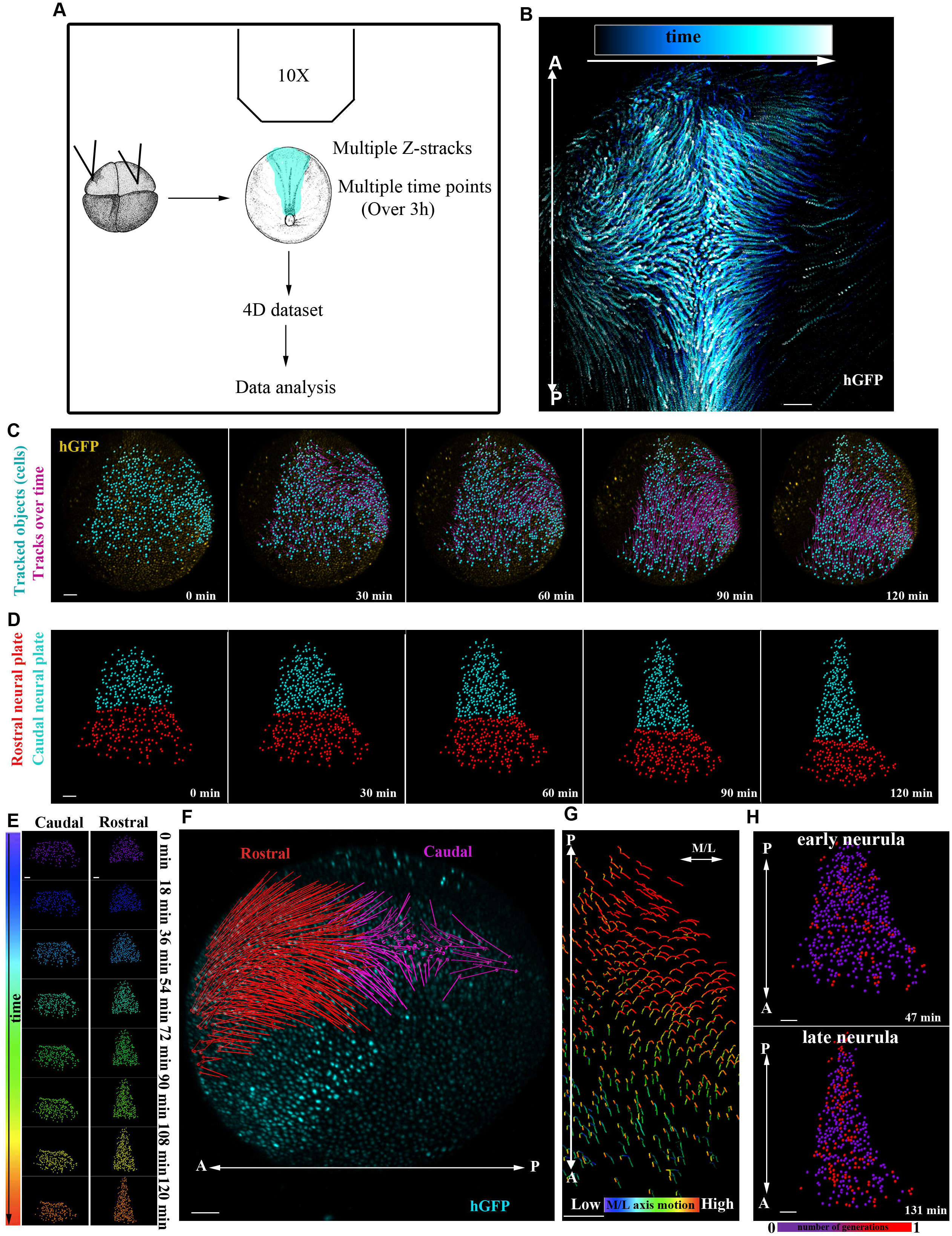
Live imaging reveals differential behaviour of caudal and rostral neural plate regions. A) Schematic representation of the method used for generation and analysis of 4D time-lapse recordings. B) Representative temporal colour coded MIP image of a time lapse recording from an embryo expressing H2B-GFP. C) Stills from a single cell tracked time lapse recording (Movie 1) of an embryo expressing H2B-GFP. D) Stills from a tracked time lapse recording revealing the behaviour of the caudal (spinal cord/hindbrain region) and rostral (midbrain/forebrain region) neural plate (NP). Caudal NP (narrows and elongates as time progress while the shape of the rostral NP remains the same. E) Temporal colour coded tracks from C displaying the evolution of rostral and caudal NP shape over time. F) Displacement map of single cell tracks overlaid over h-GFP signal at t0. The caudal NP cells move towards the midline. Rostral NP cells move anteriorly and ventrally. G) Mediolateral (ML) motion direction colour coded cell tracks revealing the absence of midline directed movement from the rostral anterior NP. H) Generation colour coded cell tracks. Proliferation is uniform within the NP during early and late neurulation. Scale bars = 100um.

### PCP mediated cell intercalative behaviour occurs only at the caudal neural plate

Polarized junction shrinkage and basal protrusive activity are both essential for PCP mediated cell intercalation during vertebrate NTC[31]. Junction shrinkage requires actomyosin contractility. Specifically, PCP signals drive the enriched localisation of actomyosin at the medial junctions within the NP [4, 17]. If the differential behaviour of anterior and posterior NP is underpinned by differential PCP driven intercalative behaviour, then the actomyosin network would be expected to display pattern variations along the AP axis of the neuroepithelium. To investigate this possibility, we stained early and mid-neurula embryos with Phalloidin to examine F-actin localisation. Subsequently, we compared the localisation of F-actin at the caudal and rostral NP. We observed that actomyosin is enriched at medial neuroepithelial cell-cell junctions only at the caudal part of the tissue, while it displayed uniform localisation at the anterior NP (**Fig 2A**). Since actomyosin localisation at mediolateral (ML) junctions is controlled by the PCP pathway, we decided to assess PCP at the posterior and anterior NP. As a readout for PCP, we used exogenously expressed Prickle2 (PK2), a PCP member previously shown to mark PCP mediated junction remodelling events within the NP[17]. As shown PK2 displayed enriched localisation at medial cell-cell junctions at the posterior regions of the NP. In contrast, PK displayed a uniform localisation at the anterior part of the tissue (**Fig. 2B**). These data are in accord with our hypothesis that differential PCP activity is behind the differential collective behaviour of the anterior and posterior NP. Junction remodelling, controlled by the PCP pathway, drives cell intercalative behaviour during NTC. Our data so far suggest that intercalative behaviour only takes place at the posterior NP. To examine this possibility, we tracked single neuroepithelial cells over time. This revealed that cells at the posterior NP display neighbour exchanges mediated by polarised T1 transitions[15]. This neighbour exchange events result in the re-orientation of the long axis of cells collectives from perpendicular to parallel to the AP axis. In contrast, anterior NP cells do not display any intercalative behaviour, with cells displaying stable contacts with their neighbours (**Fig. 2C, Movie 3**). Moreover, in agreement with the above, junction shrinkage, which takes place during polarised intercalative behaviour is evident only at the posterior part of the tissue as revealed after quantification of ML junction length over time (**Fig. 2D-E**). This polarised cell intercalative behaviour is driving CE of the posterior NP. At the same time the anterior part of the tissue does not change dimensions and simply moves to a more rostral position (**Fig. 2F, Movie 4**). In summary, the above data show that the posterior and anterior regions of the NP display distinct morphogenetic behaviour due to differential activity of the PCP pathway along the tissue AP axis.

**Figure 2.**
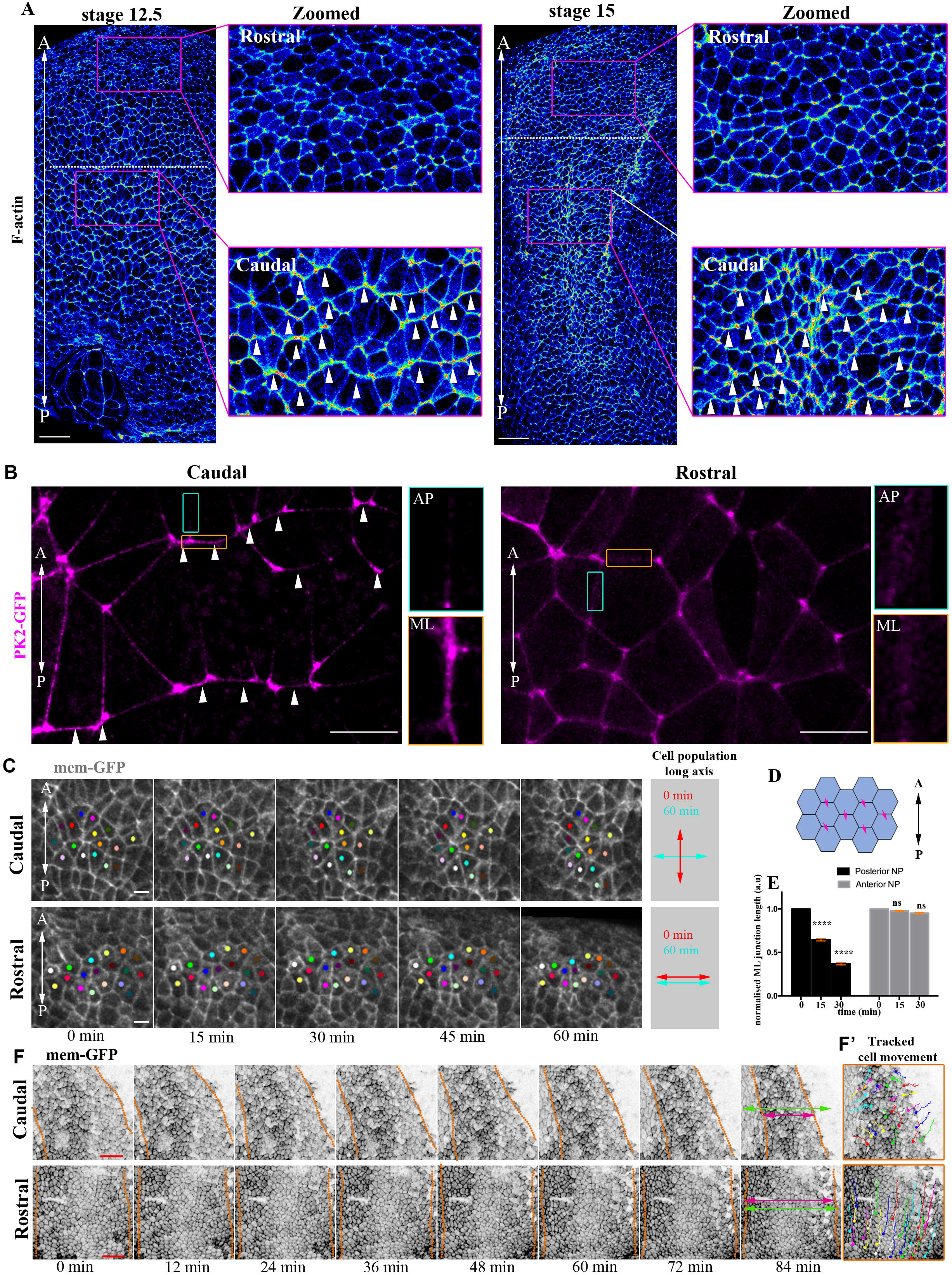
PCP mediated cell intercalative behaviour occurs only at the caudal neural plate. A) Signal intensity colour coded MIP images of representative neurula stage embryos stained with Phalloidin(F-actin). Dashed line represents the anterior/posterior neural plate (NP) limits. Arrowheads in zoomed images highlight the enrichment of F-actin at mediolateral (ML) junctions. n= 10 embryos for each stage. Scale bars = 100 um B) Representative images from a stage 14 neurula embryo exogenously expressing Prickle2-GFP (PK2-GFP). Arrowheads indicate the PK2-GFP enrichment at the ML junctions within the caudal NP. AP: anteroposterior junctions. ML: mediolateral junctions. n=5 embryos. Scale bars = 20um C) Stills from a time lapse recording of a neurula embryo. Coloured spots represent single neuroepithelial cells tracked over time. At the caudal region cells display polarized intercalative behaviour followed by neighbour exchanges. Scale bars = 20um D) Schematic showing the neuroepithelial cell-cell junctions (pink arrows) used for quantification of ML junctions’ length in (E). E) Quantification of relative ML junction length at different time points. ML junctions display reduce length as NTC progresses only at the caudal region of the embryo. Two-sided unpaired student’s t-test; ****P<0.0001; mean±SEM; n=4 embryos, 210 anterior and 210 posterior junctions F) Stills from a time lapse recording of a neurula stage embryo. Dashed lines indicate NP/SE boundaries. NP width (double headed arrows) is reduced only at the caudal NP region. Scale bars = 100um F’) Tracked movement of neuroepithelial cells at the caudal and rostral NP.

### Neural plate anterior movement requires convergent extension of the caudal neural plate region

The movement of the anterior NP towards a more rostral region takes place concomitant with the remodelling of the caudal NP due to CE. As a result of CE, the caudal NP narrows along the axis and elongates along the AP embryo axis. Therefore, the anterior NP movement might be either active or passive due to the elongation of the caudal NP. To test these possibilities, we decided to inhibit CE and then examine the movement of the anterior NP. We hypothesised that if the anterior NP moves actively its movement should not be compromised in the absence of caudal CE. In contrast if the anterior NP translocation is passive, it will be compromised if caudal CE is defective. Since CE takes place only at the posterior part of the tissue, its inhibition will allow us to specifically dissect the input of caudal CE on anterior NP movement. We decided to inhibit CE by downregulating the PCP member Vangl2, the vertebrate homologue of Vang[32]. Specifically, we used a previously characterised morpholino against Vangl2[33, 34]. To assess the effect of Vangl2 morpholino mediated downregulation we performed live imaging of Vangl2 morphant and control embryos. Comparison of NTC in control and Vangl2 morphant embryos revealed that the medial movement of the neural folds is impaired upon downregulation of Vangl2(**Fig. 3A, Movie 5**). As a result, the NP in Vangl2 morphant embryos failed to close as time progressed and remained wider than that of control embryos (**Fig. 3A**). To specifically assess if Vangl2 downregulation affected CE, we quantified medial junction shrinkage, a hallmark of polarised cell intercalative behaviour affected by Vangl2[31]. Our analysis showed that junction shrinkage was completely blocked in Vangl2 morphant embryos (**Fig. 3B**). Altogether, these results show that in Vangl2 morphants CE which takes place only at the caudal NP is impaired.

**Figure 3.**
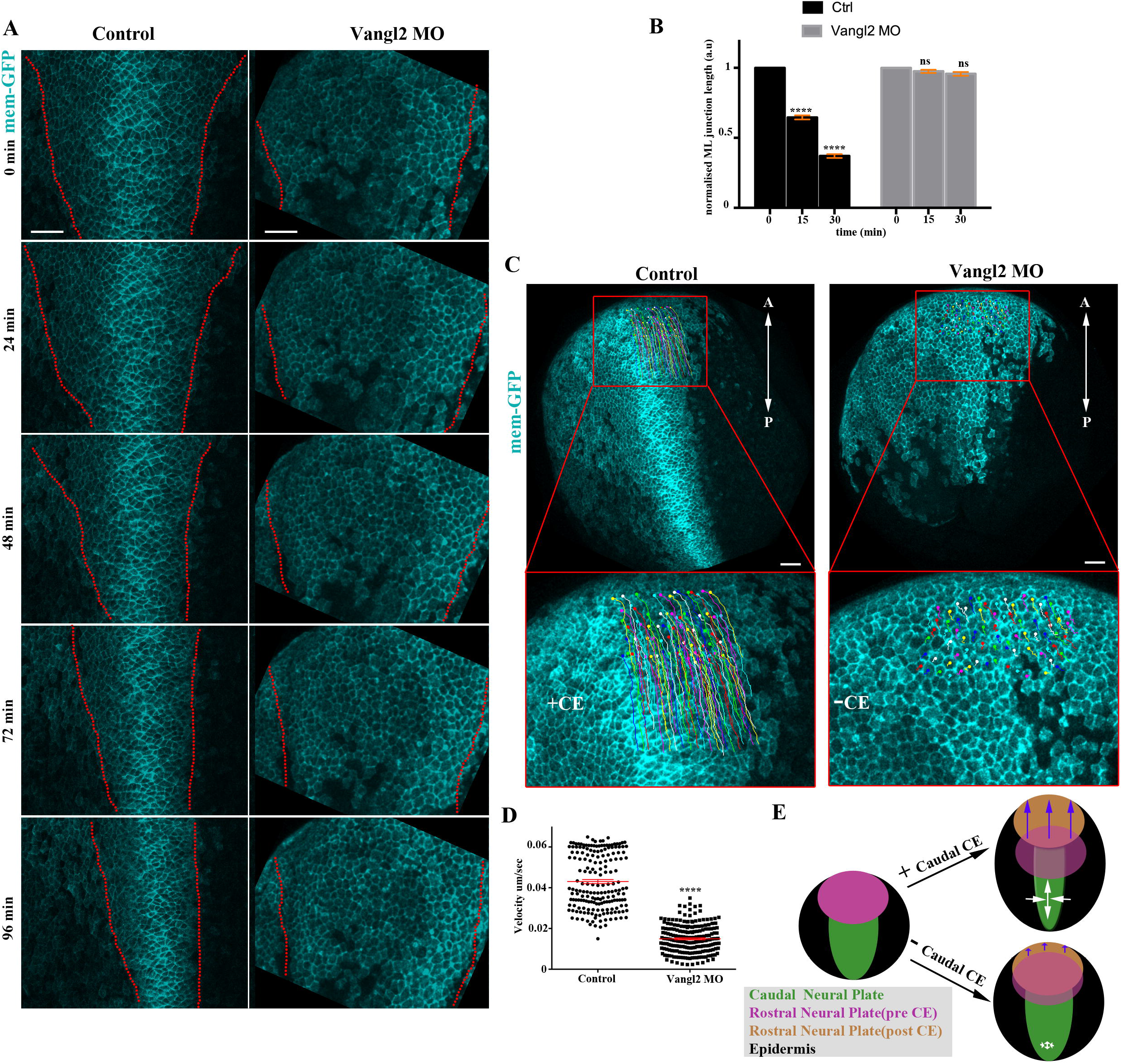
Neural plate anterior movement requires convergent extension of the caudal tissue. A) Stills from time lapse recording of representative control and Vangl2 morphant embryos. Dashed lines: neural plate (NP) boundaries. NP narrowing and neural fold movement towards the midline is defective in the Vangl2 morphant embryo. B) Quantification of mediolateral (ML) junctions’ length over time in control (4) and Vangl2 morphant(3) embryos. ML junctions display reduced length as time progress in control embryos. ML junction shrinkage is absent in Vangl2 morphants. Two-sided unpaired student’s t-test; ****P<0.0001; mean±SEM; n= 210 and 148 junctions from control and Vangl2 morphant embryos respectively. C) Representative examples of rostral neuroepithelial cell’s displacement in 60 min time period assessed by single cell tracking. The anterior directed movement of rostral neuroepithelial cells is inhibited in Vangl2 morphant embryos D) Quantification of anterior neuroepithelial cells velocity in control and Vangl2 morphant cells. Two-sided unpaired student’s t-test; ****P<0.0001; mean±SEM; n= 187 cells from 3 control embryos 209 cells from 3 Vangl2 morphant embryos. E) Model representing the contribution of caudal CE in anterior NP movement. When the caudal NP undergoes CE, its elongation leads to the anterior displacement of the rostral NP. In the absence of CE, the caudal NP does not elongate, and rostral NP displacement is absent. Scale bars = 100um.

Having validated the absence of CE in Vangl2 morphant embryos, we went on to examine how the movement of the anterior NP is affected in this context. Thus, we tracked single anterior NP cells in control and Vangl2 morphant embryos in which caudal NP CE is compromised. The anterior movement of the rostral NP was blocked in Vangl2 morphants (**Fig3. C-D, Movie 6**). This suggests that the movement of the anterior NP is passive. Specifically, CE at the caudal part of the NP results in the active narrowing and lengthening of the caudal region. The later results in the passive movement of the anterior NP towards the head region (**Fig. 3E**). In conclusion our results reveal that CE has a dual role during NTC, to drive the active remodelling of the posterior NP and to generate forces that lead to the correct positioning of the anterior part of the tissue.

### Apical constriction occurs after convergent extension and is the only contributor of anterior neural tube closure

Efficient NTC requires not only CE but also AC of neuroepithelial cells[7, 8, 35, 36]. Our results so far showed a clear contribution of CE to caudal NP morphogenesis which drives the passive movement of the anterior NP. We then went on to investigate the spatiotemporal contribution AC in relation to CE during NTC. Thus, we generated 3D time lapse recordings of embryos expressing membrane targeted fluorescent markers. This revealed that AC, which is marked by apical cell surface area reduction, follows caudal CE at the posterior part of the tissue and anterior movement of the rostral NP (**Fig. 4A-C, Suppl. Fig. 1A, Movie 7-9**). AC at the anterior part of the tissue starts simultaneously with posterior NP AC. Notably, the fusion of neural folds does not happen simultaneously along the AP axis. Neural folds first fuse at the posterior part of the tissue with the anterior most part completing the process at a later stage (**Fig. 4A-B, Suppl. Fig. 1A**). This probably happens because when AC takes place at the rostral NP the width of the spinal NP is narrower (**Fig. 4A-B, Movie 7-9**), due to the fact that CE does not take place at the anterior part of the tissue (**Fig. 4A-B, Suppl. Fig. 1 A-B, Movie 7-9**).

**Figure 4.**
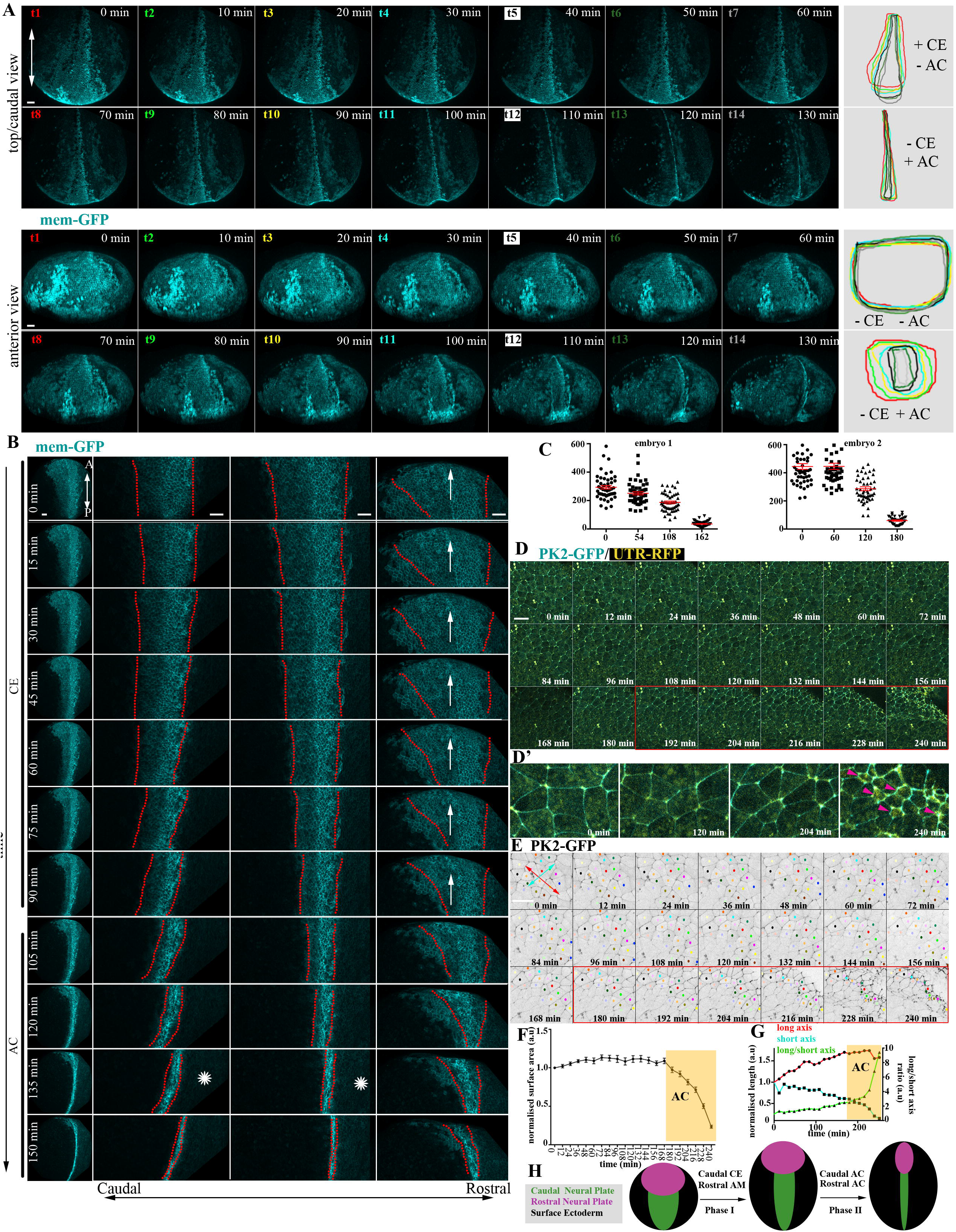
Apical constriction occurs after convergent extension and is the only contributor of neural tube closure at the rostral region. A) Stills from a 3D rendered time lapse recording of a neurula embryo. Apical constriction (AC t8-t14) occurs after convergent extension (CE t1-t7) at the caudal region of the tissue(upper panels) and at the same time occurs at the anterior part of the tissue (lower panels) which doesn’t show signs of tissue remodelling through CE. Right panels show NP shape evolution at different time points (temporal colour coded). Scale bars = 100um. B) Stills from a time lapse recording of another neurula embryo. In the left panel row, the whole embryo is shown. The next three rows show zoomed NP areas (caudal to rostral, from left to right). CE at the posterior NP precedes AC and is concomitant with the anterior displacement (arrow) of the rostral part. Posterior NTC is completed before anterior NTC (asterisk). Scale bars = 100um C) Quantification of neuroepithelial cell apical cell surface area from time lapse recordings of two different embryos. The apical cell surface area of neuroepithelial cells is decreased at 2h after imaging (imaging started at stage 13), during the last stages of NTC, showing that AC only occurs during late NTC. mean±SEM; n=50 cells for all time points except the last time point of embryo 2 which is 35 cells. D) Stills from a time lapse recording from the caudal region of a neurula stage embryo. Imaging started at stage 12.5. Red box indicates the time points when AC takes place. D’) Zoomed images of selected time points from D. The apical cell surface area of neuroepithelial cells is reduced as time progresses. Arrowheads indicate the appearance of apical-medial actomyosin during neuroepithelial cells AC. Scale bars = 50um E) Tracked zoomed region from (D). Neighbour exchanges (red and yellow cells are an example) are only present before AC takes place (red box). Double headed arrows used for quantification in (H): Red indicated the length of cell collective at the anteroposterior direction, Cyan indicates the length of cell collective at the mediolateral direction. Scale bars = 50um. F) Quantification of apical cell surface area over time. Initiation of AC is marked by the yellow box. mean±SEM: n=15 cells G) Quantification of AP length of cell collective (red line), ML length of cell collective (cyan) and the AP/ML length ratio. AP length increases through CE until the initiation of AC when it is slightly reduced indicating the absence of cell intercalative behaviour. The ML length is dramatical reduced during AC, when the tissue is bend towards the midline. H) Schematic showing the input of AC and CE during NTC. NTC occurs in two phases. During the phase I, CE of the caudal NP, drives the elongation of the caudal region of the tissue resulting in the passive anterior/ventral directed movement of the anterior tissue. During phase II, AC occurs throughout the neural plate driving the folding of the tissue and the fusion of neural folds.

To validate the observation that AC contributes to NTC after caudal CE we decided to downregulate Shroom3, a known regulator of AC[7]. Importantly, analysis of time lapse recordings of Shroom3 morphants showed that CE was unaffected in these embryos, with the midline directed movement of the neural folds unaffected (**Suppl. Fig 2A, Movie 10**). These data show that AC occurs after caudal NP CE and CE does not depend on AC. In agreement with our data, Crispr/Cas mediated knock-out of Shroom3 does not affect the polarized remodelling of posterior junctions [37].

Since, CE never takes place at the anterior part of the tissue, AC is the primary contributor of anterior NTC. Knock out mice for genes affecting AC display anterior NTDs [6, 23]. Similarly, inhibition of cell autonomous calcium flashes which regulate AC during Xenopus NTC results in defective anterior NTC (**Suppl. Fig. 2B**). Thus, while AC takes place both at the posterior and anterior NP its loss can be compensated only at the posterior part of the tissue. Thereafter, we asked if CE and AC at the posterior NP display temporal overlap. To explore this possibility, we generated high-resolution time-lapse movies of the caudal neuroepithelium. This revealed that polarised neighbour exchanges, which result in CE cease before apical cell surface area reduction begins (**Fig. 4D-G, Movie 11-12**). Therefore, we conclude that AC and CE are temporally separated during NTC. Neurulation is divided in two distinct morphogenetic phases an initial phase of CE restricted to the caudal region and a second phase of generalized AC throughout the NP. Given the lack of posterior NT closure defects in embryos with defective AC we then asked if in the absence of AC, it is possible that caudal CE might be prolonged to compensate until the two-hinge points fuse. To examine this possibility, we analysed Shroom3 morphant embryos in which AC is absent. Our analysis revealed polarised intercalative cell behaviour within the posterior NP of Shroom3 morphants, well beyond that observed in control embryos (**Suppl. Fig. 2 C-E, Movie 13**). This indicates that CE is prolonged in these embryos, moderating the effect of defective AC on NTC.

Overall, our data show that NTC is completed in two distinct phases (**Fig. 4 H**). In the first phase the caudal NP undergoes CE which drives the passive rostral movement of the anterior part of the tissue. During the second phase of NTC AC drives the bending of the tissue and the fusion of the hinge points both at the caudal and rostral regions of the NP.

### Surface ectoderm medial movement requires forces generated by neural plate morphogenesis

In addition to active force generating morphogenetic events taking place within the NP, it has been proposed that extrinsic mechanical forces also contribute actively to NTC. Both mesodermal as well as ectodermal tissues which are mechanically coupled to the NP have been suggested to act as force generators during NTC. More specifically, it has been suggested that active migration of deep SE cells is necessary for NTC in Xenopus embryos[28]. However, this is in contrast with classical studies showing that excised NPs can elongate and roll up[29]. Furthermore the argument supporting an active movement of deep epidermal cells, is inconsistent with the fact that that deep ectoderm cells from gastrula stage embryos don’t have the capacity to migrate on FN matrix when explanted from the embryo[38]. In order to examine if deep SE cells become migratory during neurulation and differ from their gastrula stage counterparts, we explanted deep SE surface cells from stage 14 and plated them on FN for 1h. Then we imaged the behaviour of explanted deep SE cells. This revealed that deep SE cells don’t acquire a migratory behaviour during neurula stages (**Fig. 5A-B, Movie 14**). To further examine the migratory capacity of the SE in the embryo we tracked deep and superficial SE cells. This revealed that deep cells move together with the superficial epithelial cells and never overtake superficial cells (**Suppl. Fig. 3A, Movie 15**), indicating that the deep SE cells don’t move autonomously but rather their movement is coupled with the movement of the superficial epithelial layer. Altogether these data suggest that deep SE cells don’t undergo active migration during NTC.

**Figure 5.**
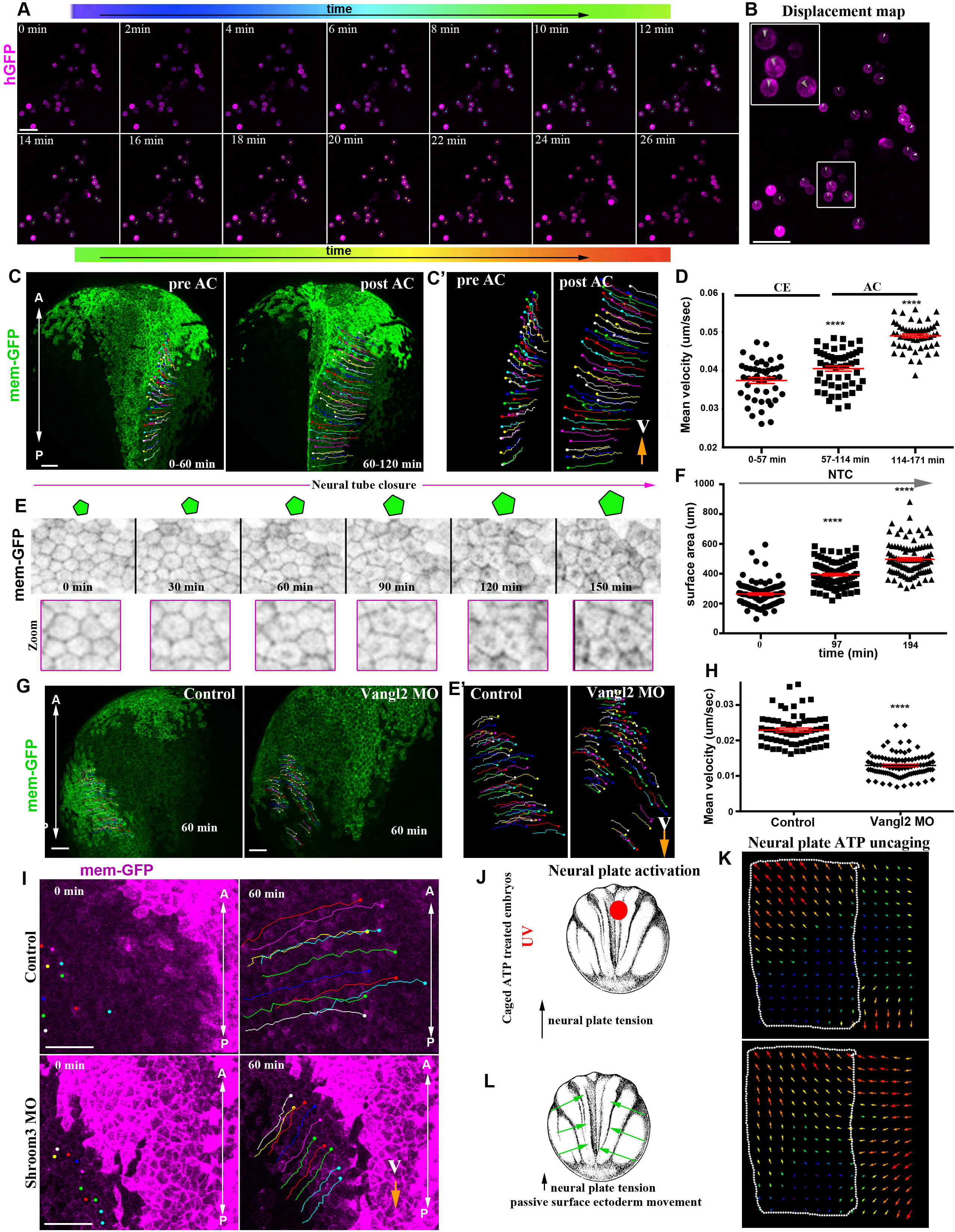
Surface ectoderm medial movement requires forces generated by neural plate morphogenesis. A) Stills from a time lapse recording of deep surface ectoderm (SE) cells plated on a FN coated coverslip. Tracks (spots) are time colour coded. B) Displacement map for a 26min time window from the time lapse recording used for cell tracking in (A) indicating absence of cell movement. C) Stills from a tracked time lapse recording of a control embryo before (left panel) and after (right panel) apical constriction (AC). C’) Zoomed tracks from C showing the increase of cell displacement and velocity (V) during AC. D) Quantification of the average cell velocity of SE cells during different phases of NTC. Two-sided unpaired student’s t-test; ****P<0.0001; mean±SEM. n=46, 53 and 54 cells for the three different time periods. E) Stills from a tracked time lapse recording showing SE cells during neural tube closure (NTC). As NTC progress, the surface area of SE cells increases. F) Quantification of the apical surface area of SE cells during neural tube closure. Two-sided unpaired student’s t-test; ****P<0.0001; mean±SEM. n=100 cells for each time point. G) Stills from a tracked time lapse recording of a control embryo (left panel) and a Vangl2 morphant embryo (right panel).G’) Zoomed tracks from C showing decrease of cell displacement and velocity (V) in the absence convergent extension in Vangl2 morphants. H) Quantification of the average cell velocity from control and Vangl2 morphant embryos. Two-sided unpaired student’s t-test; ****P<0.0001; mean±SEM, n= 63 SE cells from a control and 97 SE cells from a Vangl2 morphant embryo. I) Stills from a tracked time lapse recording of a control embryo (top panels) and a Shroom3 morphant embryo (bottom panels). SE cell’s displacement and velocity (V) is decrease in the absence of AC. J) Schematic showing the experimental approach to increase tissue tension within the NP through optogenetic uncaging of ATP. K) Particle image velocimetry (PIV) analysis illustrating increased movement speed of SE upon ATP uncaging within the NP (dashed outline). L) Schematic illustrating the behaviour of SE upon NP targeted ATP uncaging. Scale bars = 100um.

Next, we went on to examine if the SE epithelium displays an active or passive movement during neurulation. Analysis of tissue tensile properties in neurula axolotl embryos revealed that, tissue tension within the NP is higher than that of the SE[39]. Thus, the SE as a softer tissue would not be expected to generate forces able to deform the stiffer neuroepithelium. Thereafter, we formed the hypothesis that the movement of the SE towards the midline is passive as result of NT morphogenesis. To test this hypothesis, initially we tracked SE cells during NTC in control embryos. This revealed that the velocity of SE cells increases slightly during CE and shows dramatic increase when neuroepithelial cells undergo AC (**Fig. 5C-D, Movie 16**), the NTC phase with the higher rate of neural fold displacement[8]. Moreover, our single cell tracking analysis shows that the lateral SE epithelium movement mirrors the movement of the NP. The most anterior part of the SE shows an anterior directed movement while the posterior part shows a medial movement similar to the NP (**Suppl. Fig. 3B)**. Importantly quantification of cell surface area during NTC revealed that the surface area of SE cells adjacent to the NP increases as NTC progress (**Fig 5E-F**). The anterior SE, which is found beneath forebrain region, displays a unique behaviour during NTC. Specifically, this region of the SE displays an anterior and ventral directed movement during the first phase of NTC following the movement of the anterior NP which passively moves anteriorly and ventrally (**Suppl. Fig. 3C, Movie 17**). During the second phase of NTC, the cells in this region of the SE change behaviour and are stretched dorsally by the folding anterior NP and acquire an elongated shape (**Suppl. Fig. 3 D-E, Movie 18, 19**). These cell shape changes are characteristic of tissues experiencing tensile forces, generated by the active remodelling of a neighbouring tissue[40, 41].All the above findings indicate that the SE moves passively towards the midline during neurulation and its movement is coupled with NP morphogenesis.

To directly assess the later, we decided to inhibit CE and AC within the NP and then examine the movement of the SE. First, we downregulated Vangl2 within the NP by targeted injections of Vangl2 MO and we compared the movement of the SE in control and Vangl2 morphant embryos. The movement of the SE towards the midline was slower in Vangl2 morphant embryos (**Fig. 5G-H, Movie 20**), showing that NP CE contributes to the SE movement. Next, we analysed the movement of the dorsal/anterior SE during AC, in embryos where Shroom3 was downregulated within the NP. The movement of the SE was impaired in Shroom3 morphants (**Fig. 5I, Movie 21**). Overall, these data show that intrinsic NP morphogenesis, is actively pulling the SE towards the midline.

Our data so far indicate that NP morphogenesis generates intrinsic forces which pull the SE towards the midline. We decided to directly assess this interaction between the SE and the NP using optogenetics to modulate contractility in a spatially and temporally controlled manner. Specifically, we used caged ATP since ATP has been shown to be sufficient to induce cell contractility in the Xenopus ectoderm epithelium [42]. Therefore, targeted uncaging of ATP will lead to increase of cell contractility at the selected tissue region. Increase of cell contractility will subsequently lead to increased tissue tension (**Fig. 5J**). Embryos expressing membrane-GFP were treated with caged-ATP and kept in the dark until stage 14. Targeted NP ATP uncaging resulted in rapid change of SE movement direction and pace towards the excited NP region (**Fig. 5K-L, Movie 22**). This experiment shows that cells of the SE respond directly to forces generated by the NP. Overall, the above data highlight the close mechanical linkage between the NP and the SE and strongly support the notion that that SE movements during neurulation are passive and driven by force generating morphogenetic movements of the NP.

### Normal surface ectoderm homeostasis is necessary for neural tube closure

Our data highlight the close mechanical link between SE and the NP. We thus postulated that NT morphogenesis would require a balance between the active forces generated by NP morphogenesis and SE tension. Specifically, SE tension should be sufficiently low to allow tissue deformation and expansion in order to permit NTC. Integrin-β1 (Itgβ1) signalling has been shown to control cell division orientation in the ectoderm of gastrula stage embryos and absence of Itgβ1 signalling leads to tissue thickening[43]. Importantly tissue thickening has been shown to lead to increased tissue tension[44]. To test the effects of increased tissue tension on NTC we injected a previously characterized Itgβ1 MO at 1 ventral blastomere at the 4-cell stage to get embryos with unilateral SE Itgβ1 downregulation (**Fig. 6A, Suppl. Fig. 4A-B**). Itgβ1 downregulation in the SE leads to thickening of the tissue (**Fig. 6B-D, Suppl. Fig. 4C**). This is followed by impaired NTC (**Fig. 6E-F, Suppl. Fig. 4D-E, Movie 23**). This phenotype was previously attributed to the loss of active deep SE cell migration[19]. Although this interpretation cannot be ruled out it is highly unlikely to be correct based on our data indicating that SE movements are passive.

**Figure 6.**
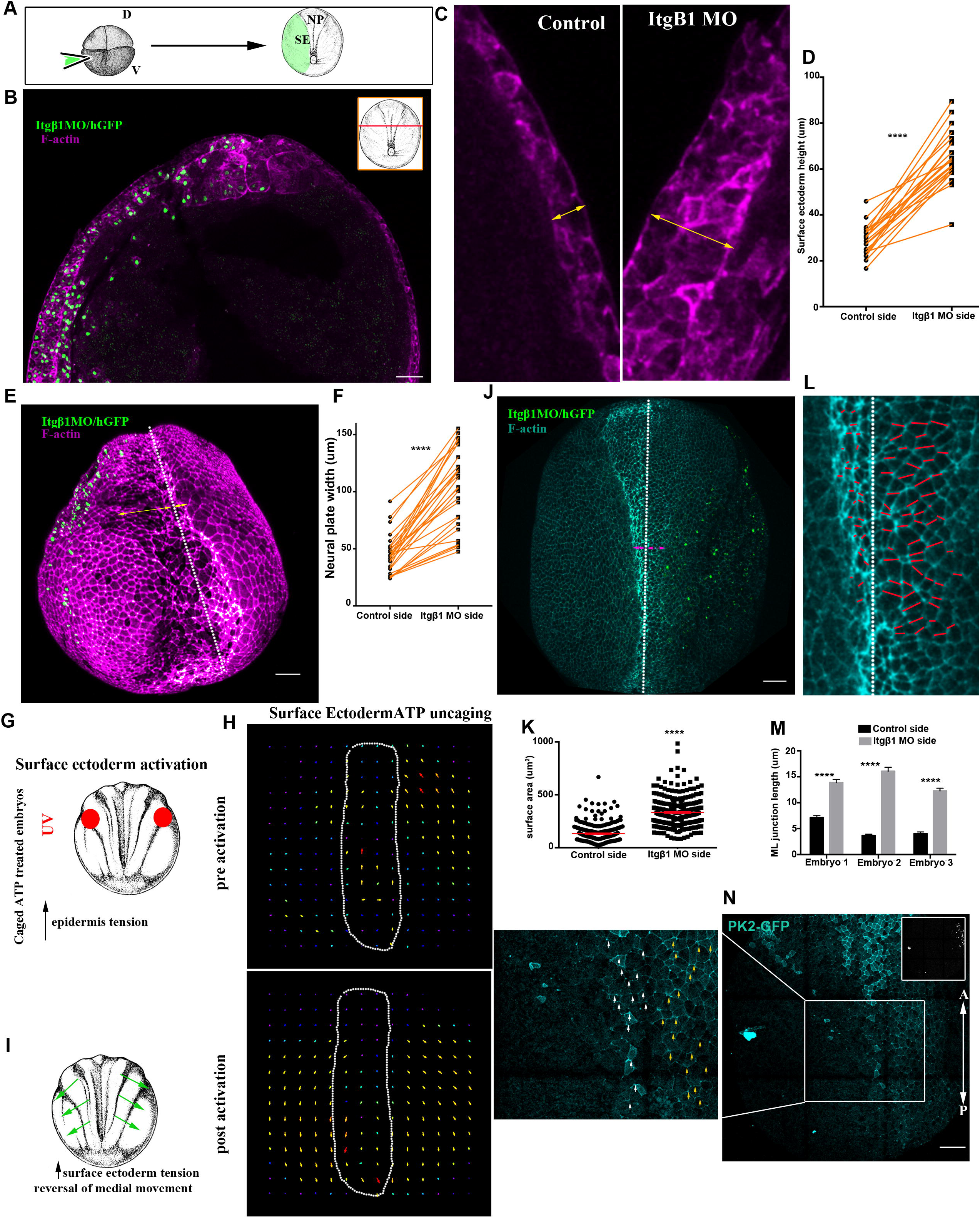
Normal surface ectoderm homeostasis is necessary for neural plate morphogenesis. A) Schematic showing the process for surface ectoderm targeted microinjections. D: dorsal. V: ventral. SE: surface ectoderm. NP: neural plate. B) Representative example from a cross section (inset) of a stage 15 embryo with SE unilateral injection of Itgβ1 MO. Itgβ1 MO was co-injected with histone-GFP. n=21 embryos. C) Zoomed images of the SE from (B). The thickness of the SE (double headed arrow) is increased at the Itgβ1 morphant side. D) Quantification of SE thickness in embryos unilaterally injected with Itgβ1 MO. Two-sided paired student’s t-test; ****P<0.0001; mean±SEM, n= 21 embryos. E) 3D image of a representative example of a stage 16 embryo unilaterally injected with Itgβ1 MO. White dashed line indicate the midline. Double headed arrows indicate the length of the NP at the control and Itgβ1 MO injected side. n=18 embryos F) Quantification of NP length at the control side and the Itgβ1 MO side in embryos unilaterally injected with Itgβ1 MO. Two-sided paired student’s t-test; ****P<0.0001; mean±SEM, n=28 embryos. G) Schematic showing the experimental approach to increase tissue tension within the SE through optogenetic uncaging of ATP. H) Particle image velocimetry (PIV) analysis illustrating reversion of SE movement upon ATP uncaging within SE during NTC. dashed outline: neural plate region. I)Schematic illustrating the behaviour of SE upon SE targeted ATP uncaging. J) Representative example of an embryo unilaterally injected with Itgβ1 MO. White dashed line indicate the midline. Double headed arrows indicate the length of the NP at the control and Itgβ1 MO injected side. n=20 embryos K) Quantification of apical cell surface area of neuroepithelial cells. The NP side with adjacent morphant SE has neuroepithelial cells with larger surface area, indicating defects in apical constriction. Two-sided unpaired student’s t-test; ****P<0.0001; mean±SEM, n=300 cells from 6 control embryos and 300 from 6 Itgβ1 morphant embryos. L) Zoomed NP from (K). White dashed line indicates the midline. Red lines indicate mediolateral (ML) junctions. ML junctions at the NP side adjacent with Itgβ1 morphant SE fail to shrink. M) Quantification of ML junction length in control NP side and NP side adjacent with Itgβ1 morphant SE. Two-sided unpaired student’s t-test; ****P<0.0001; mean±SEM, n= 50 junctions for each embryo N) Representative example of an embryo expressing PK2-GFP and unilaterally injected with Itgβ1 MO (inset H2B-GFP). PK2 displays enriched localisation at ML junctions at the control side (white arrowheads). PK2 localisation at ML junctions is unaffected at the NP adjacent with Itgβ1 morphant SE (yellow arrowheads). n=5 embryos. Scale bars = 100um.

According to our hypothesis, thickening of the SE in Itgβ1 morphant embryos would lead to an increase of tissue tension within the SE. This will affect the balance between the active forces generated by the NP and the tension within the SE leading to defective NTC. To test this hypothesis directly, we employed optogenetic induction of cell contractility. Specifically, we allowed embryos treated with caged-ATP to develop to neurula stages and subsequently uncaged ATP within the SE to increase tension in this tissue (**Fig. 6G**). We observed that upon increase of SE contractility, NTC velocity slowed down, and the direction of SE cell movement was reversed (**Fig. 6H-I, Movie 24**). These data show that defects in Itgβ1 SE morphants stem from elevated resistive forces rather than loss of deep cell migration. In summary, our data show that while the SE expands passively during NTC, its proper homeostasis and maintenance of correct tissue tension levels are permissive for NTC.

Given the NTC defects observed upon elevated tension within the SE we then asked which NP morphogenetic events are affected under elevated resistive forces from the SE. The molecular machineries controlling CE and AC are mechanosensitive. Specifically, mechanical forces can alter the distribution of PCP components which are indispensable for CE and can also affect actomyosin contractility, necessary for both AC and CE, through the control of myosin II distribution[45-47]. To specifically address how these processed are affected by elevated tension we turned to Itgβ1 unilaterally SE morphant embryos in which elevated tension is maintained throughout NTC. We initially quantified the apical cell surface area of neuroepithelial cells to assess effects on AC. This revealed that neuroepithelial cells at the side adjacent to the morphant SE had significantly larger average apical surface area (**Fig. 6 J-K**), indicative of defective AC. To assess CE, we analysed the length of ML junctions. We found that ML junctions fail to shrink at the NP side adjacent to the SE lacking Integrin signalling (**Fig. 6 L-M**) despite the fact that planar polarization of the tissue was unaffected (**Fig. 6N**). This suggests that the resistive forces from the stiffened SE cannot be overcome by the force generating machinery of the NP leading to defects in both AP and CE. Overall, our data show that while SE does not actively contribute to NT morphogenesis, its mechanical linkage to the NP imposes opposing forces thus SE morphogenesis is permissive for NTC.

## Discussion

In this study we describe the spatiotemporal contribution of the morphogenetic events taking place within the NP and we delineate the role of the SE during NTC. Our findings demonstrate that NTC is driven by a precisely orchestrated spatiotemporal pattern of morphogenetic events. Additionally, we show that NP is mechanical coupled with the SE the expansion of which is permissive for NTC.

High resolution imaging of NP morphogenesis revealed that the anterior and posterior regions of the NP display distinct morphogenetic behaviour. This is an agreement with a recent study which also detected distinct patterns of behaviour at the posterior and anterior NP [37]. Specifically, here we show that while the posterior NP narrows and elongates, the anterior part of the tissue displays a polarised anterior movement. Importantly, our data indicate that the anterior movement of the rostral NP is passive, and its correct positioning depends on the polarised intercalative behaviour of the posterior NP. The fact that we don’t observe stretching of posterior neuroepithelial cells in Vangl2 morphants excludes the possibility that anterior NP cells move actively during neurulation. If this was the case, we would expect the posterior neuroepithelial cells adjacent to the anterior NP to acquire an elongated shape as a result of tensile forces transmitted from the movement of the anterior part of the tissue.

Our results also reveal that the distinct behaviour of the anterior and posterior NP, depends on PCP mediated intercalative cell behaviour being present only at the posterior part of the tissue. Recently, it was shown that the blastopore lip found at the posterior end of the NP is a source of PCP inducing signals[48]. At the moment we cannot differentiate between the possibilities that the anterior NP can sense the signals inducing PCP and its distinct behaviour its attributed to lack of signalling possibly due to the distance from the source or if the anterior NP simply cannot respond.

CE and AC are the two major morphogenetic processes taking place within the NP. When, we analysed their spatiotemporal contribution during NTC we found that CE contributes only to the posterior NP during the first phase of NTC. During this phase the anterior NP moves towards the dorsoventral midline without any deformation. Subsequently, during the second phase AC takes place both at the posterior and anterior of the tissue with no temporal overlap with CE. In the absence of AC, CE takes place and it’s prolonged at the spinal NP, and this is likely responsible for mitigating NTC defects at the posterior NP when AC is defective. The distinct spatiotemporal contribution of AC and CE during NTC can explain the differences in the phenotypes described upon loss of function of genes regulating CE vs AC. Mutations resulting in NTDs most of the times affect either the anterior or the posterior part of the tissue but rarely both. Specifically, mutations in genes regulating AC have been shown to induce anterior NTDs while genes regulating CE have been shown to induce posterior NTDs. Our results suggest that mutations in genes contributing to AC don’t induce posterior NTDs because in the absence of AC the posterior tissue continues to move towards the midline through prolonged CE.

The contribution of extrinsic forces, generated by neighbouring tissues, to NP morphogenesis has been controversial throughout the years. Here we focused on the role of the SE on NTC. Our results demonstrate that the deep SE cells don’t have a migratory capacity in contrast with a previous study[28]. These results are in agreement both with the fact that deep ectoderm cells from gastrula embryos are not migratory[38], and with the fact that SE explants developed up to tailbud stages never display any migratory behaviour. Specifically, SE explanted on a FN substrate always remain round and never spread like their mesoderm counterparts suggesting that both superficial and deep layers are passive. Using live imaging we now show that the SE movement mirrors the movement of the NP, and its movement is directly influenced by NP morphogenesis. We also show that induction of contractility within the NP through optogenetics, is sufficient to elicit directed movement of the SE towards the NP midline. Finally, we show that deep and outer cells move synchronously and in the same direction. Overall, our results show that the movement of the SE towards the midline is passive and is dependent on forces generated by NP morphogenesis. Similar mechanically coupled morphogenesis between neighbouring tissues a has been described during early mouse development[49], drosophila development[40] and chick embryo axis elongation[50].

In addition, to our findings that the SE is mechanically coupled with the SE, in this study we examined how defective SE development might affect NTC. We found that thickening of the SE epithelium, after loss of Itgβ1 signalling leads to NTDs. It has been suggested previously that Itgβ1 signalling is necessary for the migration of the deep ectodermal cells towards the midline[28]. Here we ruled out this possibility since we showed that the deep SE cells are not migratory. Thickening of the ectoderm in Xenopus embryos has been shown to result in tissue stiffening[44]. To examine the effect of SE tissue stiffening we optogenetically activated cell contractility within the SE. This blocks the movement of the NP hinge points pointing out that a balance between the forces generated by the NP and the tension within the SE ensures proper NP development. Opposing forces by the surrounding tissues have been documented during NTC in mouse embryos[51]. Our data directly show that increased tension within the SE prevents its deformation in response to NP generated pulling forces and as a result mechanically opposes NTC. Similarly, overexpression of Grhl2 in mice results in the increase of tissue tension and defective NTC[45, 52].

Last, we show that defective SE development affects both AC and CE within the NP. At the same time PCP remains unaffected. This can be explained by the fact the molecular regulators of actomyosin which executes both AC and junction shrinkage during CE have been shown to be mechanosensitive[46, 47]. It is likely that resistive forces stemming from the SE are higher than the capacity of actomyosin force generation by either CE or AC. In agreement with this, a recent report has demonstrated that ectopic induction of AC can block AC in a neighbouring cell population that is genetically programmed to undergo AC[53]. Thus, our results demonstrate the necessity of proper SE development for NTC and highlight the close mechanical linkage between the two tissues and the mechanosensitive nature of NP morphogenesis.

## Materials and Methods

### Xenopus embryos and microinjections

Female adult *Xenopus laevis* frogs were induced to ovulate by injection of human chorionic gonadotropin. Eggs were fertilised *in vitro*, after acquisition of testes from male frogs, dejellied in 2% cysteine (*pH* 7.8) and subsequently reared in 0.1× Marc’s Modified Ringers (MMR). mRNA and microinjection. For microinjections, embryos were placed in a solution of 4% Ficoll in 0.33× MMR and injected using a glass capillary pulled needle, forceps, a Singer Instruments MK1 micromanipulator and Harvard Apparatus pressure injector at the 4 cell stage according to Neiuwkoop and Faber[54]. After injections, embryos were reared for 1 hour in 4% Ficoll in 0.33× MMR and then washed and maintained in 0.1× MMR. Injected embryos were allowed to develop to neurula stage (Nieuwkoop and Faber stage 12-5–13) at 17°C and imaged live, or allowed to develop to the appropriate stage and the dissected or fixed in 1x MEMFA [55] for 1-2 hours at room temperature. Capped mRNAs encoding fluorescent protein fusions were *in vitro* transcribed using mMessage machine kits (Ambion). The amount of mRNA per 4nl of microinjection volume was as follows: membrane-GFP, 100 pg; histone-RFP, 80 pg; Prickle2-GFP, 100 pg; Utrophin-RFP, 80pg.

### Morpholino oligonucleotides

The Shroom3[7], Vangl2[33] and Itgβ1[28] morpholinos(MO) were previously described and were ordered from GeneTools. 30ng of Shroom3 MO, 20ng of Vangl2 MO, 30ng Itgβ1 MO were injected per blastomere. Shroom3 and Vangl2 MOs were injected at dorsal blastomeres of 4 cell stage embryos to target the neural plate. Itgβ1 MO was injected at the ventral blastomere of 4 cell stage embryos to target the surface ectoderm.

### Immunofluorescence

Immmunofluorescence was performed as previously described[56]. Briefly, embryos were fixed for 2 hours at room temperature, permeabilized in PBST (1 × PBS, 0.5% Triton, 1% dimethyl sulfoxide) and blocked for 1 hour in 10% donkey serum. Primary antibodies were incubated overnight at 4 °C. We used a primary antibody against integrin β1 (1:50, 8C8, Hybridoma Bank). Embryos were washed in PBST and incubated for 2 h with secondary antibodies at RT, washed several times and post-fixed in 1XMEMFA. Secondary antibodies used were Alexa fluor 488 (1:500, Invitrogen), Phalloidin was incubated together with the secondary antibodies, phalloidin 546(1:500, Invitrogen).

### Deep surface ectoderm migration assay

Explants of deep cell surface ectoderm were dissected from stage 13 embryos. Dissections were performed in 1× MBSH (88 mM NaCl, 1 mM KCl, 2.4 mM NaHCO_3_, 0.82 mM MgSO_4_, 0.33 mM Ca(NO_3_)_2_, 0.33 mM CaCl_2_, 10 mM Hepes, and 10 μg/ml Streptomycin and Penicillin, pH 7.4). Single cells were dissociated in alkaline buffer (88 mM NaCl, 1 mM KCl, and 10 mM NaHCO_3_, pH = 9.5) as previously described[38]. Dissociated cells were plated on FN-coated glass bottom dishes and left to adhere for 45 and then imaged.

### Nifedipine treatment

For nifedipine treatment, 500× stock of nifedipine (Sigma) in DMSO were added to the medium at stage 13 at 200 μM. Embryos were fixed at stage 25.

### ATP uncaging

DMNPE-caged ATP (A1049; Molecular Probes) was added to the medium to a concentration of 100 μM 1hour before imaging. For the photolysis of DMNPE-caged ATP, the samples were illuminated for 45s with UV light. The duration of illumination was controlled manually

### Live imaging

Live imaging of neurula stage Xenopus embryos was performed on a ZEISS LSM 710 confocal microscope with a Plan-Apo 10X, NA 0.45 objective for all time lapse recordings except for Figure 4D-E, Movie 11 and 12 for which we used a Plan-Apo 40X, NA 1.1 objective. The ZEISS ZEN software was used during imaging. Embryos were imaged in a custom chamber made of thick layer of vacuum grease on a microscope slide and sealed with a coverslip. Embryos were mounted in 0.1x MMR and kept at room temperature during imaging.

### Image analysis and single cell tracking

All image analysis and quantification were carried out using Fiji software[57]. For single cell tracking (Figure 1), the spots function of the Imaris image analysis software was used. Automated generated tracks were manually curated. Single cell tracking in Figures 2F,3C, 5C, 5G, 5I and Suppl. Fig. 3C was carried out using the manual tracking plugin of Fiji.

### Statistics

GraphPad Prism 6.0 software was used for all statistical analysis performed. The sample size of the experiments carried out was defined based on previous experimental experience. Quantitative data presented, shows the mean ± s.e.m., or the total number of datapoints obtained. The statistical tests carried out on the quantitative data obtained are annotated in each figure legend.

## Author Contributions

N.C conceived the project, designed and carried out the experiments and data analysis. N.C and P.A.S wrote the manuscript.

## Competing Interests

The authors declare no competing interests.

## Acknowledgments

We thank Dr. John Wallingford, for kindly providing the PK2-GFP plasmid.

## Funding

This work was co-funded by the European Regional Development Fund and the Republic of Cyprus through the Research and Innovation Foundation (Project: POST-DOC/0718/0087).

## Notes

### Competing Interest Statement

The authors have declared no competing interest.

