## Supplementary figures and images for "Distinct spatiotemporal contribution of morphogenetic events and mechanical tissue coupling during Xenopus neural tube closure"

### Supplementary Figure 1

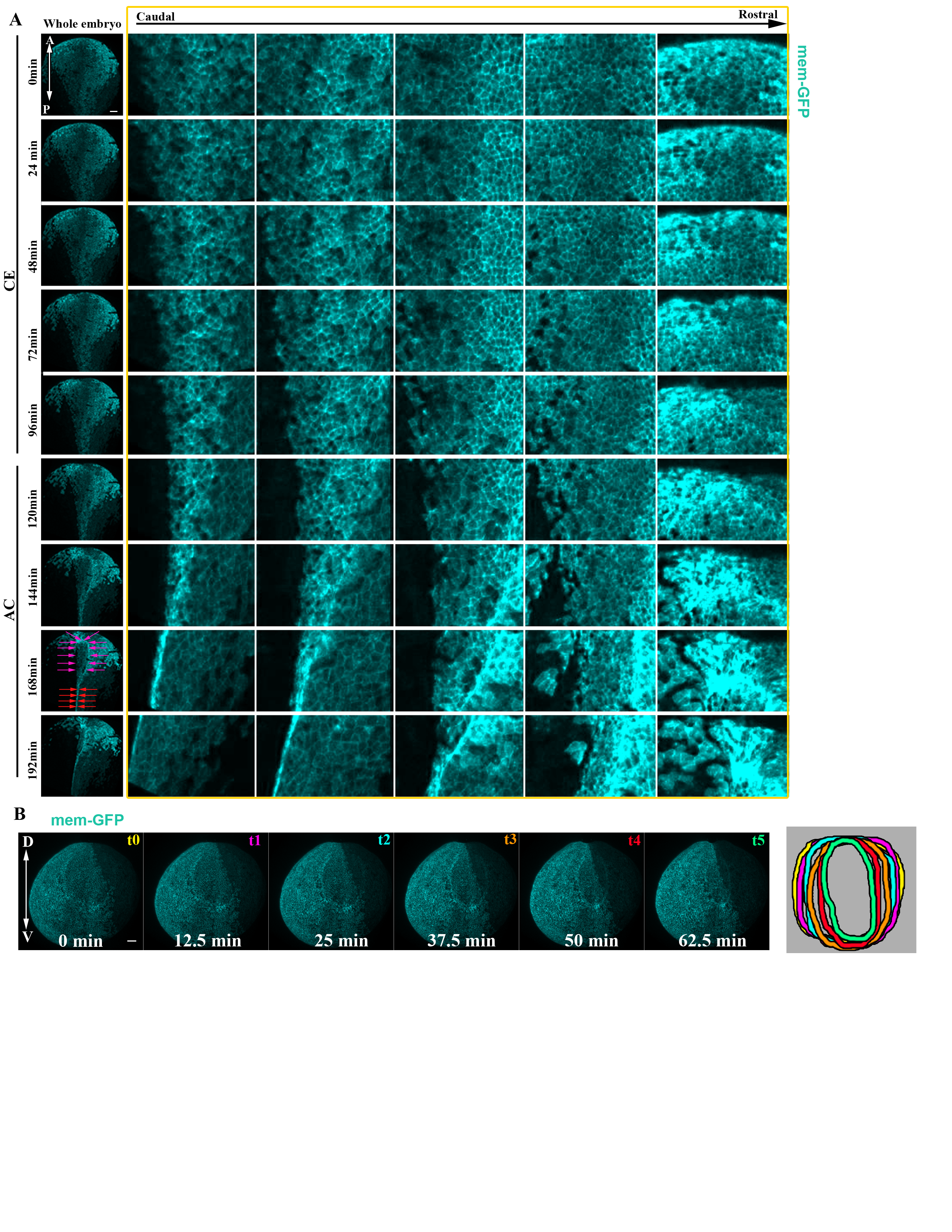

### Supplementary Figure 2

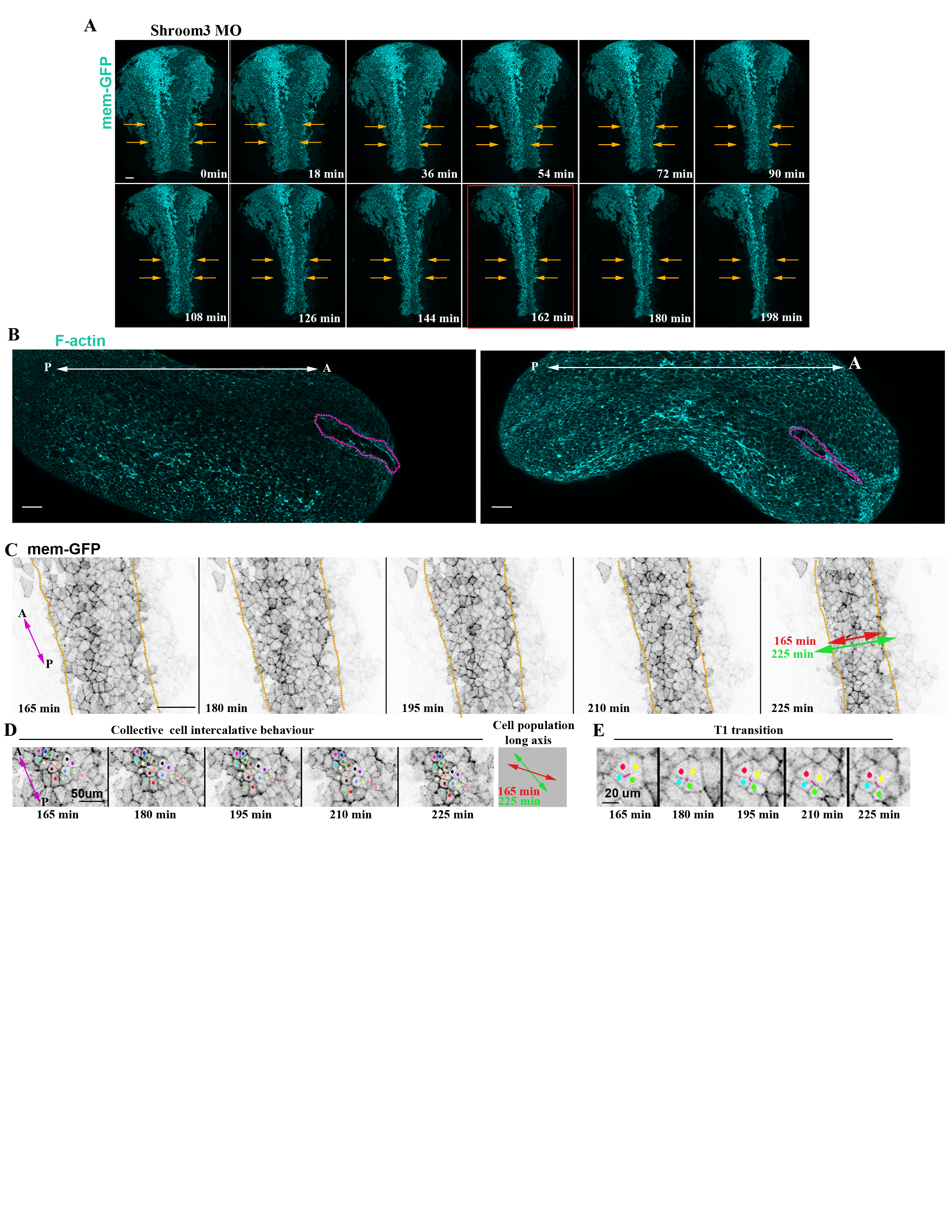

### Supplementary Figure 3

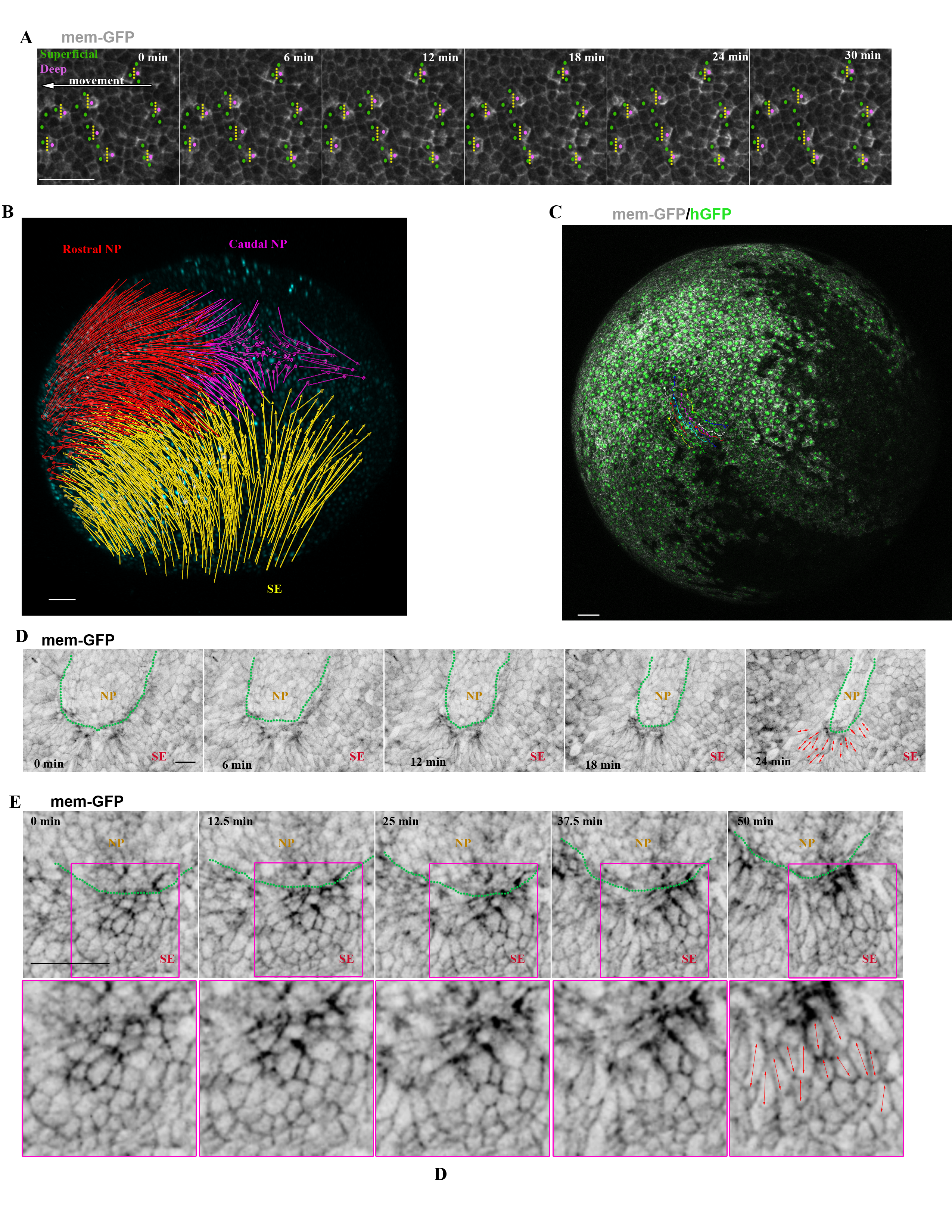

### Supplementary Figure 4

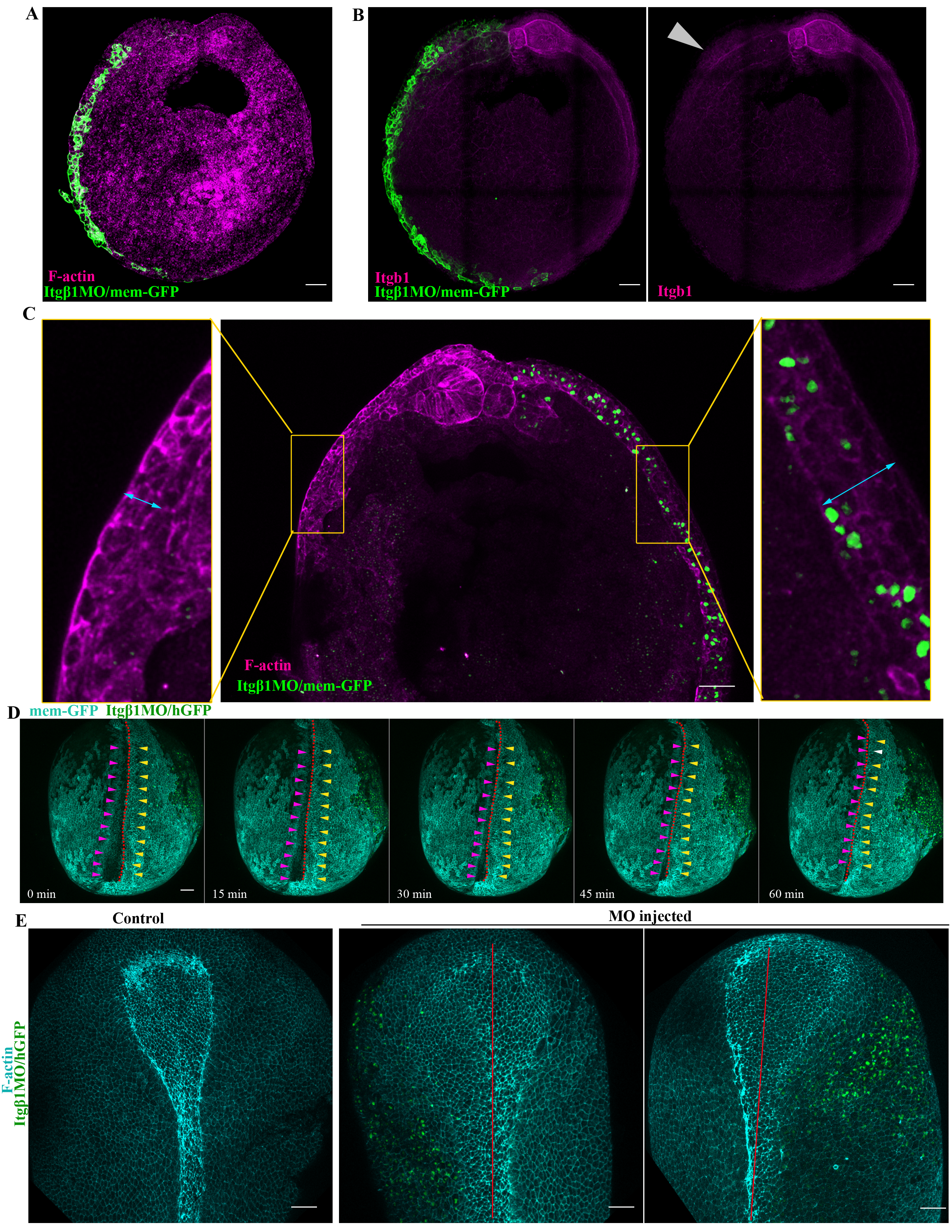
