## Supplemental legends for "Distinct spatiotemporal contribution of morphogenetic events and mechanical tissue coupling during Xenopus neural tube closure"

**Supplementary Figure legends**

**Supplementary Figure 1 related to Figure 4.** A) Stills from a time lapse recording of a neurula embryo. In the left panel row, the whole embryo is shown. The next five rows show zoomed NP areas (caudal to rostral from left to right). CE at the posterior NP precedes AC. Caudal NTC (red arrows) is completed before anterior NTC (magenta arrows). B) Stills from a time lapse recording of an embryo expressing membrane-GFP. Right panel shows the shape (temporal colour coded) of the rostral neural plate during NTC indicating absence of CE. The anterior NP shrinks along the ML axis due to AC. A: anterior. P: posterior. D: dorsal. V: ventral. Scale bars = 100um.

**Supplementary Figure 2 related to Figure 4.** Stills from a time lapse recording of a Shroom3 morphant embryo. The caudal neural plate narrows (arrows) narrows and lengthens through CE. By the time-point that control embryos have completed NTC (red box), NTC is not completed, and caudal NP continues to narrow . B) Representative embryos (stage 25) treated with 200uM nifedipine from stage 14. The anterior neural tube failed to close (dashed outline). n=20 embryos. C) Zoomed stills from the same time lapse recording as in (A), starting from the time point highlighted with red box in (A), when NTC is completed in control embryos. Dashed lines: neural plate boundaries. NP CE continues well beyond the time point of NTC completion in control embryos. Double headed arrows: NP width at different time points. D)Singe cell behaviour analysis from a zoomed region from (C) showing polarised cell intercalative behaviour and neighbour exchanges (red and orange marked cell as an example). Right panel: The cell population long axis reorients as time progress, becoming parallel with the embryo’s AP axis. E) An example of T1 transition from (C). The remodelled cell junction is highlighted with magenta. Scale bars for A, B and C = 100um, for D= 50um and for E=20um.

**Supplementary Figure 3 related to Figure 5. A)** Stills from a time lapse recording of a control embryo expressing showing the surface ectoderm (SE). Superficial SE cells are marked with green spots and deep SE cells with magenta dots. Note that as NTC progress the deep SE cells move together with superficial cells and never overtake superficial cells. B) Displacement map of single cell tracks coloured according to their spatial location. The movement of SE cells mirrors the movement of neuroepithelial cells. C) Representative example of rostral/ventral SE cell’s displacement in a 60 min time window during the first phase of NTC, assessed by single cell tracking. The cells move towards the rostral and ventral side of the embryo. D) Stills from a time lapse recording focusing on the anterior neural plate (dashed line). During anterior NP folding (second phase of NTC) the SE cells beneath the NP are stretched and acquire an elongated shape (double headed arrows). E) Stills from a time lapse recording of a distinct embryo focusing on the anterior neural plate (dashed line). During anterior NP folding the SE cells beneath the NP are stretched and acquire an elongated shape (double headed arrows). Scale bars = 100um.

**Supplementary Figure 4 related to Figure 6.** A) Cross section of a representative embryo. Microinjection of Itgβ1 MO+ membrane GFP at the animal side of 1 ventral blastomere at the 4-cell stage led to unilateral SE targeting. B) Cross section of a representative embryo unilaterally targeted with Itgβ1 MO at the SE showing effective downregulation of Itgβ1 at the MO injected side (arrowhead). C) Representative example from a cross section of an embryo with SE unilateral injection of Itgβ1 MO. Itgβ1 MO was co-injected with histone-GFP. Zoomed images of the SE reveal that the thickness of the SE (double headed arrow) is increased at the Itgβ1 morphant side. D) Stills from a time lapse recording of a unilateral SE Itgβ1 morphant embryo. NTC at the side adjacent to the Itgβ1 morphant SE is defective. Arrowheads indicate the neural folds (magenta: control; yellow: Itgβ1 morphant side). The neural fold at the Itgβ1 morphant side fail to reach the midline (dashed red line) E) Representative examples of a control embryo and two unilateral Itgβ1 SE morphant embryos. Red line indicates the midline. Scale bars = 100um.

**Supplementary Movie legends.**

**Movie 1:** Single cell tracked time lapse recording of a neurula stage embryo expressing histone-GFP. In the first part of the video all neuroepithelial cells tracks are shown with cyan spots and magenta lines. During the second part of the video the neuroepithelial cells are colour coded according to their spatial location within the neural plate. Cyan dots: caudal neural plate. Red dots: rostral neural plate. Time interval = 3 min.

**Movie 2:** Generation colour coded cell tracks of neuroepithelial cells during neural tube closure. Proliferation is uniform within the neural plate during neurulation. Time interval = 3 min.

**Movie 3:** Distinct behaviour of neuroepithelial cells at the caudal (left) and rostral (right) regions of the neural plate of an embryo expressing mem-GFP. Neuroepithelial cells display polarised intercalative behaviour only at the caudal part of the tissue. Time interval = 3min.

**Movie 4:** Part 1: Caudal (left) and rostral (right) morphogenesis neural plate morphogenesis from an embryo expressing mem-GFP. Only the caudal neural plate undergoes convergent extension Part 2: Single cell tracking of neuroepithelial cells reveals the presence of cell intercalation at the caudal neural plate (left) and an anterior directed movement of the rostral part of the tissue (right). Time interval= 3 min.

**Movie 5:** Time lapse recording of a control (left) and a Vangl2 morphant embryo. Neural plate CE is defective in Vangl2 morphant embryo. Time interval= 3min

**Movie 6:** Tracking of rostral neuroepithelial cells in control (left) and Vangl2 morphant embryos (2 right embryos). The anterior directed movement of rostral neuroepithelial cells is absent in Vangl2 morphant embryos. Time interval= 3min

**Movie 7:** First example of neural tube closure (NTC). Neural tube closure at the caudal (top) and rostral neural plate(bottom). Initially, caudal neural plate undergoes CE and rostral neural plate moves towards the dorsoventral midline. Subsequently AC occurs both at the caudal and anterior neural plate. Caudal NTC is completed before rostral NTC. Time interval= 3min

**Movie 8:** Second example of NTC**.** Initially, caudal neural plate undergoes CE and rostral neural plate moves towards the dorsoventral midline. Subsequently AC occurs both at the caudal and anterior neural plate. Caudal NTC is completed before rostral NTC. Time interval= 3min

**Movie 9.** Third example of NTC. Initially, caudal neural plate undergoes CE and rostral neural plate moves towards the dorsoventral midline. Subsequently AC occurs both at the caudal and anterior neural plate. Caudal NTC is completed before rostral NTC. Time interval= 3min

**Movie 10.** Time lapse recording of a Shroom3 morphant embryo. Caudal neural plate CE is not affected. Time interval= 3min

**Movie 11.** Time lapse recording showing cell behaviour at the caudal neural plate from stage 12.5-stage 16. Initially neuroepithelial cells undergo polarised cell intercalation. During the last phase of neural tube closure, neuroepithelial cells undergo apical constriction, marked by the reduction of their apical cell surface area and enrichment of medio-apical actin signal (yellow). CE and AC do not show temporal overlap. Cyan: Prickle2, Yellow: Utr-GFP. Time interval= 30 sec.

**Movie 12.** Manual single cell tracking of a zoomed region from Movie 11 using the PK2-GFP signal. Neighbour exchanges are present during CE but not during AC. Time interval= 12.5 min.

**Movie 13.** Time lapse recording showing a zoomed region of a Shroom3 morphant neural plate from movie 10. This time lapse recording starts for the time-point during which NTC is normally completed in control embryos. CE continues at the caudal NP of Shroom3 morphant at time points when CE does not take place in control embryos. Single cell tracks showing the collective cell intercalative behaviour (left) and an example of a T1 transition (right). Time interval= 3min

**Movie 14.** Time lapse recording of deep surface ectoderm (SE) cells expressing hRFP and plated on a FN coated coverslip.

**Movie 15.** Time lapse recording showing the behaviour of the surface ectoderm during neural tube closure. SE moves towards the medial direction (left). Deep surface ectoderm cells (magenta) never overtake superficial SE cells (green). Time interval = 3 min.

**Movie 16.** Single cell tracking of surface ectoderm cells before (left) and after(right) neural plate AC. Time interval= 3 min

**Movie 17.** Single cell tracking of anterior/ventral SE cells during the first phase of NTC. Time interval= 3 min.

**Movie 18.** First example of anterior/ventral SE cell behaviour during the second phase of NTC. Time interval= 20 sec.

**Movie 19.** Second example of anterior/ventral SE cell behaviour during the second phase of NTC**.** Time interval= 150 sec.

**Movie 20.** SE movement in control (left) and Vangl2 morphant embryo (right). Neural plate midline is on the right side. Time interval= 3 min.

**Movie 21.** Tracked SE movement in control (left) and Shroom3 morphant embryo (right). Neural plate midline is on the right side. Time interval= 3 min.

**Movie 22.** Time lapse recording showing the behaviour of the neuroepithelium (left) and the surface ectoderm (right) before and after ATP uncaging within the neural plate upon UV excitation (bright area).

**Movie 23.** Neural tube close in an embryo with uniliteral SE Itgb1 downregulation. NTC at the NP side adjacent to the morphant SE is defective. Red overlay. highlights the defective NTC at the affected side. Time interval= 6 min

**Movie 24.** Time lapse recording showing the behaviour of the neuroepithelium (middle) and the surface ectoderm (right and left) before and after ATP uncaging within the surface ectoderm upon UV excitation (bright areas).
